# Antibiotic Resistance Gene Variant Sequencing is Necessary to Reveal the Complex Dynamics of Immigration from Sewers to Activated Sludge

**DOI:** 10.1101/2023.03.05.531174

**Authors:** Claire Gibson, Susanne A. Kraemer, Natalia Klimova, Bing Guo, Dominic Frigon

## Abstract

Microbial community composition has increasingly emerged as a key determinant of antibiotic resistance gene (ARG) content. However, in activated sludge wastewater treatment plants (AS-WWTPs), a comprehensive understanding of the microbial community assembly process and its impact on the persistence of antimicrobial resistance (AMR) remains elusive. An important part of this process is the immigration dynamics (or community coalescence) between the influent and activated sludge. While the influent wastewater contains a plethora of ARGs, the persistence of a given ARG depends initially on the immigration success of the carrying population, and the possible horizontal transfer to indigenously resident populations of the WWTP.

The current study utilised controlled manipulative experiments that decoupled the influent wastewater composition from the influent microbial populations to reveal the fundamental mechanisms involved in ARG immigration between sewers and AS-WWTP. A novel multiplexed amplicon sequencing approach was used to track different ARG sequence variants across the immigration interface, and droplet digital PCR was used to quantify the impact of immigration on the abundance of the targeted ARGs. Immigration caused an increase in the abundance of over 70 % of the quantified ARGs. However, monitoring of ARG sequence variants at the immigration interface revealed various immigration patterns such as (i) suppression of the indigenous mixed liquor variant by the immigrant, or conversely (ii) complete immigration failure of the influent variant. These immigration profiles are reported for the first time here and highlight the crucial information that can be gained using our novel multiplex amplicon sequencing techniques. Future studies aiming to reduce AMR in WWTPs should consider the impact of influent immigration in process optimisation and design.

## 1 Introduction

Antimicrobial resistance (AMR) is recognized as one of the greatest threats to public health worldwide (Abadii et al., 2019). Each year 700,000 deaths are attributed to AMR and without action this number is predicted to rise to 10 million by 2050 (O’Neill, 2014). The United Nations Environment Assembly (UNEA-3) have recognised the importance of the environment in the development, spread and transmission of AMR to humans and animals (United Nations Environment Programme (UNEP), 2022). Of particular interest are wastewater treatment plants (WWTPs), which have been identified as hotspots of AMR (Rizzo et al., 2013) and gateways to the environmental spread. Although a reduction in the load is observed, the wastewater treatment process does not effectively remove all phylogenetically mobile antibiotic resistance genes (ARGs) and resistant bacteria (ARB) before release into the environment (Lapara et al., 2011). Consequently, effluent wastewater has been shown to contribute to antibiotic resistance in surface waters and sediments downstream of effluent discharge points (Quintela-Baluja et al., 2019; Reichert et al., 2021). ARGs are also disseminated in the waste biosolids produced during the treatment process (Munir et al., 2011; Gao et al., 2012), which are often applied to agricultural land as fertilisers which creates another route of AMR dissemination. To minimize the spread of AMR, wastewater treatment plant design requires urgent optimization for the removal of ARB and ARGs.

With increasing knowledge of emerging contaminants in wastewater and the benefits of water resource recovery, the need for new wastewater treatment technologies is widely recognised. In this context, studies have aimed to minimise ARG release into the environment with novel design. However results are often variable. For example, some studies of anoxic-aerobic membrane bioreactor observed a reduction in the abundance of ARGs (Le et al., 2018; Zhu et al., 2018), whilst others found the abundance of genes such as *tet*C to increase (Xia et al., 2012). The use of ozonation to reduce ARG loads resulted in removal efficiencies which varied between ARG classes (Staley et al., 2019). Similarly, some found chlorination to cause large reductions in the abundance of ARGs (Zhang et al., 2019) and increases in the abundance of cell free ARGs (Liu et al., 2018), whilst others reported no impact on genes such as *bla*TEM-1 (Pang et al., 2016). These contradictory results exemplify our lack of understanding of the drivers in the persistence and proliferation of ARBs and ARGs in WWTPs, which remains one of the greatest hurdles in developing appropriate treatment strategies to reduce AMR.

Influent wastewater contains a plethora of ARB which harbour several ARGs with specific sequence variants often associated with the genetic context of the gene (Gibson et al., 2023b). Each ARG sequence variant is likely to obey different elimination or persistence mechanisms following their immigration into the biological treatment process from the sewer. The characterization of these mechanisms, however, requires the identification and quantification of ARG variants across the influent and activated sludge (AS-WWTP) interface with high resolutions and sensitivity (Smith et al., 2022; Gibson et al., 2023b). Although quantitative PCR is sensitive, it is uninformative with regards to ARG sequence variants. Conversely, metagenomic shotgun sequencing can provide information on the genetic context of the most abundant variants, but it has limited capabilities in the identification of variants occurring in low abundance, and it is 10^2^ to 10^5^ less sensitive than qPCR for ARG detection. Numerous studies have used data on ARG occurrence in the influent and activated sludge to infer the origin of AMR. However, more recent studies demonstrate that ARGs can vary at the sequence level based upon their origin (Zhang et al., 2021). The development of specific approaches to address this knowledge gap is the main goal of the current study.

The fate of immigrating ARB and ARGs is related to the complex ecological dynamics occurring at the interface between the influent and AS-WWTP. Firstly, the persistence of a given ARG variant depends on the immigration and success of its carrying population in the downstream community (Gibson et al., 2023a). Some ARGs may persist because the carrying population is already an indigenous resident of the AS-WWTP community. In this case, the ARG variant may not even be present in the influent community. Secondly, the persistence of a given ARG variant may also be impacted by its phylogenetic mobility. Horizontal gene transfer has been demonstrated to play a pivotal role in the spread of ARG across species (Von Wintersdorff et al., 2016). Although a given ARB may be unsuccessful in the AS-WWTP community, processes such as conjugation, transformation and transduction enable the movement of ARGs and increase the likelihood of ARG persistence by transferring to other, better adapted, hosts. For example, conjugative plasmids have been shown to play a significant role in facilitating the persistence of multidrug resistant ARB in WWTPs (Che et al., 2019). To determine the contributions of these mechanisms, ARG sequence variants need to be tracked across the immigration interface between the influent wastewater and the activated sludge. The current study employed a novel multiplex amplicon sequencing approach (Smith et al., 2022; Gibson et al., 2023b) to study ARG sequence variant dynamics with the required sensitivity and resolution. Such a technique will likely provide valuable insights into the contradictory results on the persistence of ARGs.

The compounded impact of complex immigration dynamics, community composition drifts and horizontal gene transfers on the fate of ARGs are best studied in highly controlled and replicated reactor experiments. In full-scale WWTP, the influent substrate compositions, bacterial population, and ARGs vary on a daily basis (Guo et al., 2019; Sun et al., 2021). These variations, even when small, limit our ability to accurately assess the exact mechanisms behind influent immigration and AMR in AS-WWTPs. Therefore, this study utilised triplicated lab-scale reactors fed with a synthetic wastewater, which allowed for a high level of control on the immigration process and the possibility of reproducing similar conditions in subsequent manipulative experiments. Among the operated reactors, the only varying factor was immigration, which was simulated by the addition of naturally occurring microbial communities (i.e., suspended solids) harvested in municipal wastewaters to the synthetic wastewater.

The ARG analyses presented here expand upon previous observations on the population dynamics and community assembly obtained with the same experimental set-up and published previously (Gibson et al., 2023a). Droplet digital PCR was used to quantify the variation in abundance of the ARGs across the immigration interface. Due to influent wastewater containing ARGs originating from several sources (e.g., clinical waste, sewer biofilms and agricultural runoff (Guo et al., 2019; Qin et al., 2020)), it was hypothesised that ARGs variants from the influent wastewater would be different than the ones occurring endogenously in the activate sludge. Thus, multiplex amplicon sequencing was used for ARG source tracking.

## 2 Methods

### 2.1 Samples

Samples for analysis were collected from reactors previously operated as described by Gibson et al. (Gibson et al., 2023). Briefly, small scale activated sludge (AS) reactors were inoculated with mixed liquor taken from three full scale AS-WWTPs (Figure 1: Block A-La Prairie AS, Block B-Cowansville AS and Block C-Pincourt AS). Reactors were operated with a hydraulic retention time of 1.8 days and a solid retention time of 5 days, and fed with a synthetic wastewater (Syntho; Boeije et al., 1999) to ensure a stable wastewater composition over time. During Phase 1 of reactor operation, all reactors received Syntho only to ensure that a stable community was formed and any existing immigrants were removed. In Phase 2, to investigate the impact of immigration, in one third of the reactors the synthetic wastewater was supplemented with influent solids (i.e., an influent wastewater community) taken from a full-scale AS-WWTP. Two control groups were also included, the first was a substrate control, that received Syntho supplemented with sterile influent solids (autoclaved at 121°C, 15 psi for 30 minutes). The final 9 reactors received Syntho only to act as a continuity control. Finally, during Phase 3 of reactor operation, influent solids were removed from the feeds to determine if the impact of immigration was maintained over time.

**Figure 1.**
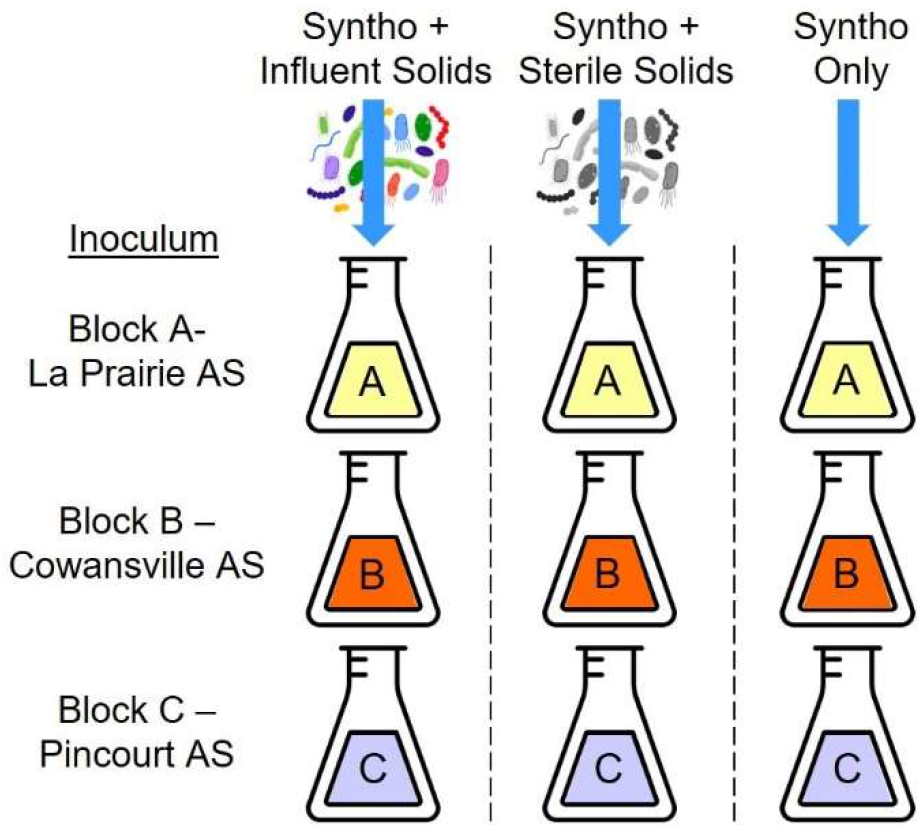
One set of Reactors which received one source of influent solids (27 reactors). Reactors were inoculated with three different activated sludge communities (Block A-C). One third of the reactors received synthetic wastewater (Syntho) supplemented with influent solid which was the group with immigration (9 reactors). Another third of the reactors received Syntho and sterile solids which acted as a substrate control (9 reactors). The final third of reactors received Syntho only and acted as a continuity control (9 reactors).

This experiment was repeated with the use of three different sources of influent wastewater community. It was observed that the impact of immigration was similar in each set of reactors, thus only one set of reactors was selected for ARG analysis (27 reactors). The chosen reactors received influent solids from Cowansville AS-WWTP (Set B; Gibson et al., 2023). For ARG analysis, samples from each Block (with the same inoculum; Block A-La Prairie AS, Block B-Cowansville AS and Block C-Pincourt AS), receiving the same feed were pooled (3 biological replicates pooled). A total of 12 samples collected during Phase 2 of reactor operation were analysed: 3 groups with immigration, 3 groups with sterilised influent solids, 3 with synthetic wastewater only, and 3 influent wastewater samples as shown in Figure 1.

### 2.2 Sample Collection and Processing

Mixed liquor biomass samples were collected from the reactors and centrifuged in 2 mL microcentrifuge tubes for 5 mins at 16,000 × g. The supernatant was discarded and biomass was stored immediately at −80 °C until nucleic acid extraction (approximately 3 months). Three aliquots were stored per reactor sample. DNA was extracted from approximately 0.25 grams of each stored biomass sample using DNeasy PowerSoil Kit (Qiagen, Germantown, MD, USA) following the manufacturers protocol with a final elution volume of 100 μL. One blank was included per batch of extractions (approximately 24 samples). Samples were extracted in duplicate, and DNA aliquots were immediately stored at −80 °C until future use.

The quality of nucleic acids was assessed using the ratio of absorbance at 260 nm and 280 nm (260/280) obtained using NanoDrop™ One. DNA extracts with a 260/280 value between 1.8-2.0 were considered to have good purity. Quant-iT™ PicoGreen™ dsDNA Assay kit (Invitrogen, Ontario) was used to accurately quantify the double-stranded DNA concentration of the extracts prior to droplet digital PCR.

### 2.3 Droplet Digital PCR

A total of 15 different antibiotic resistance genes were analysed using droplet digital PCR and multiplexed amplicon sequencing (Table S1). Clinically relevant ARGs were selected which displayed resistance to five classes of antimicrobials (beta-lactams, fluoroquinolones, macrolides, tetracyclines) to investigate whether the impact of immigration varied with ARG class. In addition, four multidrug resistance genes were included. Primers were designed by Swift Biosciences, Ann Arbor and manufactured by Integrated DNA Technologies, USA. Appropriate DNA dilutions were determined using a standard quantitative PCR reaction (Powerup SYBR green master mix) performed using three DNA dilutions to check for PCR inhibition. The fold dilution for digital droplet PCR was calculated using *Eq*. 1. Samples were then diluted to the appropriate concentration using DNase/RNase free water.

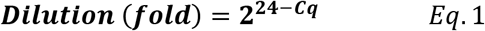

A 20 μL reaction mix was prepared for each droplet digital PCR reaction, which included 1 x QX200™ ddPCR™ Evagreen® Supermix (Bio-Rad, CA, USA; catalogue number 1864033), 100 nM of the forward and reverse primers, template DNA as determine using equation 1, and RNase/DNase-free water. For quality control purposes, a positive and no template control were included in each run. Droplets were generated using the Bio-Rad QX200™ Droplet Generator and DG8™ Cartridges with 20 μL of reaction mix and 70 μL of Droplet Generation Oil for Evagreen (Bio-Rad, CA, USA). Following droplet generation, droplets were transferred to a 96 well plate and sealed with foil using a Bio-Rad PX1 PCR Plate Sealer. A Bio-Rad C100 Touch Thermal Cycler was used for thermal cycling of all plates. Reaction conditions used were as follows; 95 °C for 5 min., followed by 50 cycles of 95 °C for 30 sec., *x* °C for 60 sec. and 72 °C for 30 sec. whereby *x* is the annealing temperature (Table S1). Followed by 4 °C for 5 min., 90 °C for 5 min. and a hold at 12 °C. Annealing temperatures were optimised using a gradient PCR method on a pooled DNA sample. The optimal annealing temperature was selected based upon the separation between positive and negative droplets, and the lowest ‘rain’ i.e. droplets defined as neither positive or negative. The primer concentration was optimised by conducting a test assay on a representative pooled DNA sample, at various primer concentrations ranging from 100 – 250 nM. A final primer concentration of 100 nm was selected for the assays.

Droplets were analysed using the Biorad QX200™ Droplet Reader and the absolute quantification (ABS) experimental setup. Results were visualised using the QuantaSoft™ Software (Bio-Rad, version 1.7.4) to provide an absolute quantification of the ARG per nanogram of DNA. Reactions producing fewer than 10,000 droplets were excluded and the droplet digital PCR reaction was repeated. The threshold was defined using the auto-select function in the QuantaSoft™ software (Bio-Rad, version 1.7.4). In cases where an appropriate threshold was not automatically selected, a threshold was manually chosen. All samples were analysed in duplicate in different PCR runs to confirm the technical reproducibility. Copies of each ARG/ng-DNA were normalised to the copies of 16S rRNA gene/ng-DNA to provide a relative ARG copy number/16S rRNA gene.

### 2.4 Sequence Diversity of ARGs

A multiplexed amplicon sequencing approach was used to analyse sequence variant diversity among the ARGs detected in the reactors. ARGs were amplified using a multiplex PCR kit (Qiagen) with the following reaction conditions; 95 °C for 15 min. followed by 25 cycles of 94 °C for 30 sec., 60 °C for 90 sec. and 72 °C for 60 sec., and a final extension step at 60 °C for 30 min. Amplicons obtained from individual PCR reactions (using one set of primers per reaction) were pooled and used as a positive control. The products of the multiplex PCR reaction were purified using SPRI Select beads (Beckman Coulter) to remove primer dimers at a sample to bead ratio of 0.8. Samples were barcoded and pooled at equimolar concentration. The pooled samples were sequenced on the Illumina MiSeq PE250 platform at McGill University and Génome Québec Innovation Centre (Montréal, QC, Canada).

### 2.5 Bioinformatics

Reads from each sample were split according to their forward and reverse primer sequences allowing for one sequencing error per primer sequence using a custom script available upon request. Read pairs where both or one of the reads was missing a primer sequence at the beginning or where the primer sequences found did not correspond to each other were removed. Subsequently, we used Trimmomatic v. 0.39 to trim all reads using the following settings: LEADING: 3 TRAILING: 3, SLIDINGWINDOW:4:15 and MINLEN:36. Then we used vsearch v2.13.3 to merge the trimmed read pairs before converting them to fasta format. The merged reads where then sorted and clustered with an identity of 1 and a minimum length of 100 bases while keeping track of the cluster size using the - sizeout option.

PCR and sequencing errors created many unique or very low count sequence variants that we removed using a custom R script. First, we tested if the distribution of read counts for each sequence variant produced by an individual primer pair followed a Poisson distribution (centered around the mean of all counts) and corrected the resulting p-values (subtracted from 1 such that sequence variants with high read counts had low p-values) using the Bonferroni correction. If none of the sequence variant have a corrected p-value below 0.5, we removed the whole dataset, as this indicated that we are not able to distinguish real sequence variants from those generated by PCR or sequencing errors (noise sequence variants). If any sequence variants survived the filtering, we aimed to remove the noise sequence variants from the datasets by finding the minimum of a density function based on the total data for each gene (including noise sequence variants) and removing all sequence variants with counts less than the minimum found. By comparing with other filtering approaches (e.g., DADA2 (Callahan et al., 2016) or minimum absolute value), it was found that the filtering adopted here was the most stringent, but was still able to recover the diversity of synthetic mixes of variants.

We subsequently combined all reads corresponding to a specific resistance gene (including multiple primers amplifying the same gene). Files were re-replicated using a custom python script available upon request (using the Pandas module), before constructing a table of allelic distribution across samples equivalent to an OTU table for each resistance gene using vsearch v2.13.3’s cluster_fast option with the otuout option. Unique consensus sequence variants were obtained using thecentroids option.

To ensure that the resulting sequence variants were the products of amplification of the target ARG, we blasted all consensus sequences to a custom ARG database using the diamond (v0.9.32.133) blastx option with an e-value of 3 and a minimum identity of 80. Consensus sequences that were not similar to any entries in the ARG database were removed from the analysis.

### 2.6 Statistics

The Mann-Whitney U test was used to determine if there was a statistically significant increase in the abundance of each ARG with immigration. Samples taken from the reactors with immigration (receiving Syntho and influent solids) were compared to the control groups at a significance level of p<0.05.

Procrustes analysis was performed using the R Vegan package (Oksanen et al., 2022) to determine whether the microbial community of the reactors correlated with the ARG profile. Procrustes was used to compare the ARG and microbial community ordinations. ARG data was standardised to means of 0 and standard deviation of 1. Using the R Vegan package, Principal Component Analysis (PCA) was conducted for the ARG dataset based on the Euclidean distances. Principle coordinate analysis (PCoA) was performed using Jaccard distance on the microbial community data, as this distance was previously shown to most effectively display the impact of immigration on the community in the current experiment (Gibson et al., 2023). The function “Protest” was used with 999 permutations to test the significance between the configurations.

## 3 Results

### 3.1 Impact of immigration on the abundance of ARGs in the activated sludge

Reactors were operated under highly controlled conditions to test the impact of immigration on antibiotic resistance in the activated sludge. All 27 reactors in the three Blocks (3 different inoculum) received a synthetic wastewater (Syntho) feed, which allowed the influent wastewater composition to be carefully controlled over time. To ensure the reactors were operating at steady state, the chemical oxygen demand and suspended solids concentrations were monitored over time. After operating the reactors for 12 SRTs and ensuring steady state was reached, during Phase 2 in each Block (9 reactors in total; Block A-La Prairie AS, Block B-Cowansville AS and Block C-Pincourt AS), three reactors received synthetic wastewater with added influent solids to simulate the impact of immigration. Another three of the reactors received autoclaved influent solids and acted as a substrate control. The final three reactors received Syntho only as a continuity control to establish a baseline for the level of AMR without immigration.

The impact of influent immigration on the activated sludge microbial community is discussed at length in Gibson et al., 2023. Briefly, immigration impacted the microbial community composition of the activated sludge, and reactors with immigration became more similar to the microbial community of the influent solids. Up to 25 % of sequencing reads were observed to be contributed through influent immigration, representing a significant proportion of the activated sludge community. Using a mass balance approach, it was observed that the growing immigrant population typically exhibited a lower and often negative net growth rates in the activated sludge, when compared to the core resident genera which typically displayed a positive net growth rate. In Gibson et al., 2023, focus was placed on the impact of immigration on the microbial community composition of the activated sludge alone, whilst in this publication the impact on AMR is explored.

To determine the impact of immigration on AMR in the AS, digital droplet PCR was used to quantitatively assess the concentration of fifteen ARGs in the reactor samples and influent wastewater. Results showed that immigration caused a significant increase in the abundance of eleven of the fifteen ARGs in the activated sludge when compared to the control groups (sterile control and no immigration control together) (Figure 2; Mann Whitney U Test; p<0.05). Genes observed to increase in abundance were distributed in several classes of antimicrobial resistance such as ARGs against fluoroquinolones (*qnr*B and *qnrS*), beta-lactams (*bla*TEM and *bla*MOX), macrolides (*dfr*A and *mph*E), and tetracyclines (*tet*Q and *tet*O). Potentially alarmingly, three out of four of the efflux pump-associated multi-antimicrobial resistance genes quantified showed a significant increase in concentration with immigration (*mar*R, *msr*D and *rob*A). Genes such as *bla*MOX were present in the reactors with and without immigration, whilst others such as *mar*R and *qnr*B were detected in the activated sludge mixed liquor only with immigration. Therefore, immigration not only impacted the concentrations of specific ARGs, but also the diversity of ARGs detected.

**Figure 2.**
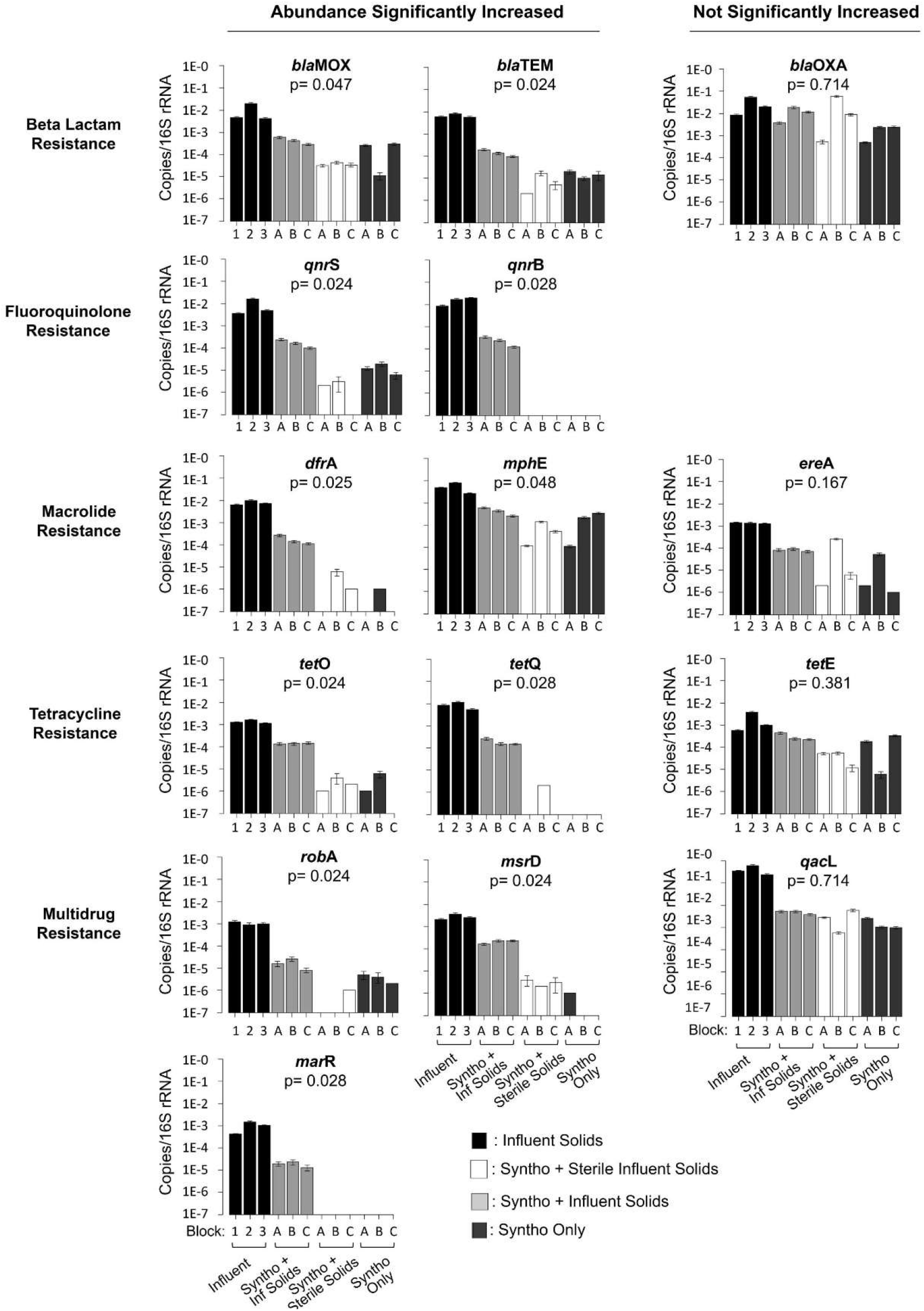
Quantification of antibiotic resistance genes using droplet digital PCR. Reactors with immigration received Syntho and influent solids. Reactors receiving Syntho and sterile influent solids acted as a substrate control. Reactors receiving Syntho only acted as a no immigration continuity control. The Block represents the inoculum source of the reactors; A-La Prairie ML, B-Cowansville ML, C-Pincourt ML, Influent samples are numbered based upon the order received by the reactors during Phase 2. Each bar represents a composite sample of three biological replicates. The Error bars represent the standard error between duplicates. The Mann-Whitney U Test was used to assess the statistical significance of the change in abundance of each ARG with immigration when compared to the sterile and no immigration control combined.

Whilst the concentrations of selected ARGs increased with immigration, others such as *bla*_OXA_ and *qac*L did not significantly change (Figure 2). The presence of these genes in similar abundance under all reactor conditions suggests that they are likely associated with the core resident populations, which were defined as taxa that occurred under all reactor conditions (i.e., independent of immigration).

Overall, the relative abundance of numerous ARGs in the AS significantly increased with immigration. Despite this increase, the relative abundance of ARGs (copies/16S rRNA) in the reactors typically remained lower than in the influent wastewater stream. However, it should be noted that the mixed liquor contained a higher concentration of biomass, and consequently the absolute ARG concentrations likely differed. The absolute ARG concentrations (Table S2) were calculated using information on reactor operation such as the volatile suspended solid concentration, yield of DNA/g-VSS, 16S rRNA gene/ng-DNA and gene abundance (16S rRNA gene) (Table S4, Gibson et al., 2023). The greatest reduction in the absolute concentration between the influent and mixed liquor was observed for the *rob*A gene which reduced by 1.14 logs of gene copies/L. Other genes such as *bla*OXA displayed an increase of 0.42 log gene copies/L between the influent and mixed liquor. Consequently, despite the reduction in ARG relative concentrations (copies/16S rRNA gene) between the influent solids, high absolute ARG concentrations remained within the activated sludge.

### 3.2 Correlation between the microbial community composition and ARG profile

In numerous studies, the microbial community composition has increasingly emerged as a key determinant of ARG content (Forsberg et al., 2014; Wu et al., 2017; Zhou et al., 2017). In continuity with these observations, a Procrustes analysis was performed to determine whether the observed changes in the abundance of ARGs in the reactors correlated with changes in the microbial community composition with immigration. The previous analysis of the community composition data of the current reactor experiment demonstrated that the impact of immigration was better visualised using the Jaccard dissimilarity than the Bray-Curtis dissimilarity, suggesting that immigration impacts taxa at lower abundances in the mixed liquor (Gibson et al., 2023). Similarly, Procrustes analysis using the Jaccard distance matrix revealed a significant correlation (Procrustes M^2^ = 0.60, *p* = 0.015) between the microbial community compositions of the reactor activated sludge and the profiles and concentrations of ARGs they carried (Figure 3b). However, this correlation was not significant (Procrustes M^2^ = 0.73, *p* = 0.290) when using the Bray-Curtis dissimilarity (Figure 3d). Taken together, this suggests that the changes in the ARG profiles observed is due to immigration of taxa mainly occurring at low abundances within the activated sludge.

**Figure 3.**
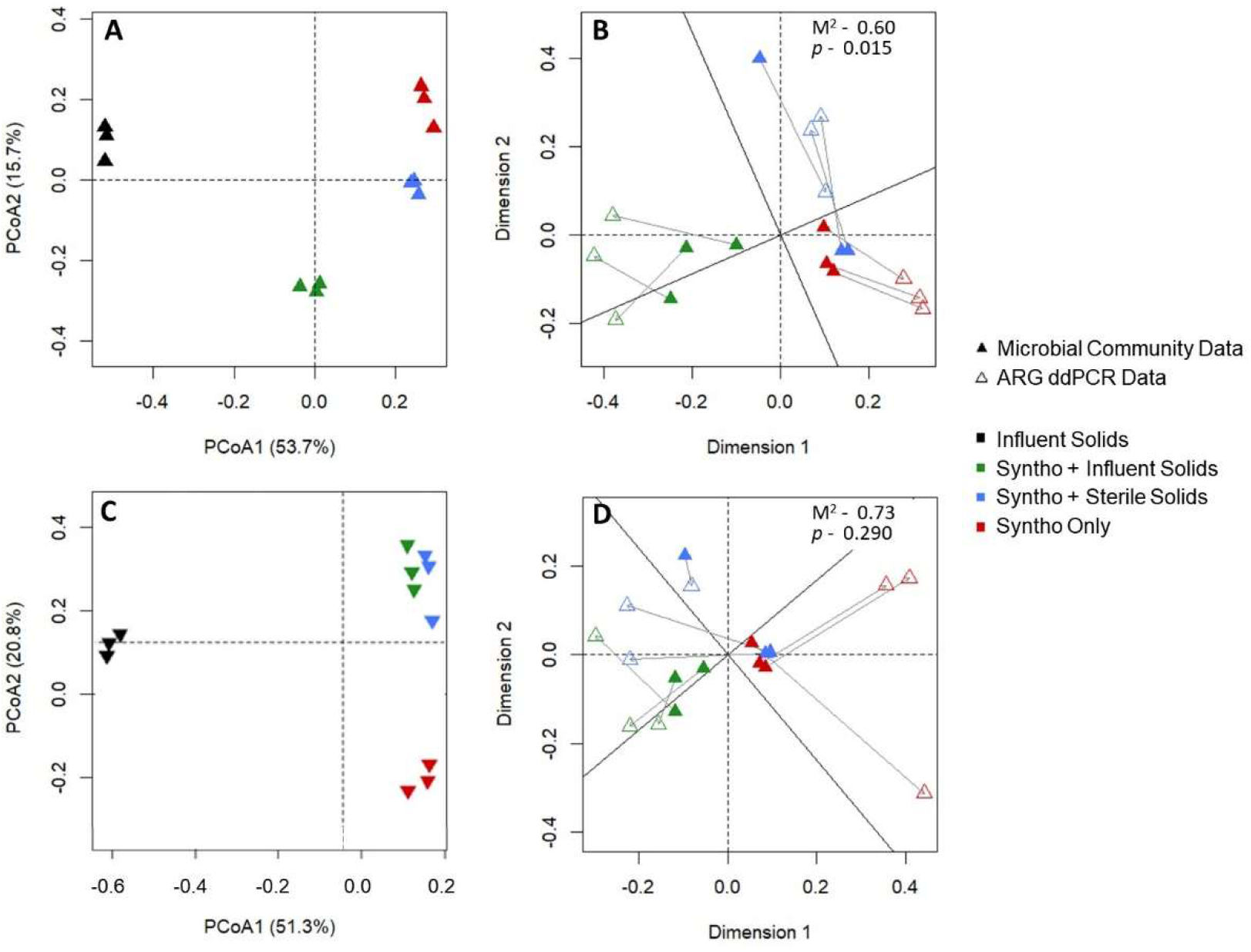
Procrustes Analysis to investigate the correlation between the microbial community and ARG composition. a) Microbial Community of the reactors visualised using Jaccard dissimilarity as determined by 16S rRNA gene sequencing b) Procrustes analysis of microbial communtiy and ARG profile with Jaccard dissimilarity c) Microbial Community of the reactors visualised using Bray Curtis dissimilarity as determined by 16S rRNA gene sequencing d) Procrustes analysis of microbial communtiy and ARG profile with Bray Curtis distance.

### 3.3 Analysis of ARG Sequence Variants

Digital droplet PCR showed that over 70 % (11 out of 15) of the ARGs investigated significantly increased in abundance with immigration. However, questions remained about the exact dynamics occurring at the interface between the influent and activated sludge. Chief among them was whether the increase in abundance of a relative ARG was due to variants originating from the influent. In other words, were the observed changes in the activated sludge likely due to direct ARG immigration or other changes induced by the presence of influent solids? To investigate the immigration dynamics at greater depth, a multiplexed amplicon sequencing approach was used to detect ARG sequence variants within the influent and reactor samples.

After stringent variant filtering based on the Poisson distribution, sequence variant information was obtained for eleven of the fifteen ARGs analysed (Figure 4). The four remaining targets (*msr*D, *mar*R, *qnr*B and *qnrS*) were undetected after the filtering process either because they did not survive the stringent filtering process (*msr*D and *qnr*B), or insufficient reads were obtained resulting in no detection (*qnr*S and *mar*R). Amplicon sequencing of the eleven ARGs revealed different variant distributions or concurrence profiles between the influent and the activated sludge mixed liquors (Figure 4). In several cases, the ARG sequence variants were specific to either the influent or reactors, with few examples of concurrence in the reactors and the received influent solids. This demonstrates that PCR based methods over-simplified the ARG dynamics at the interface between the influent and activated sludge.

**Figure 4.**
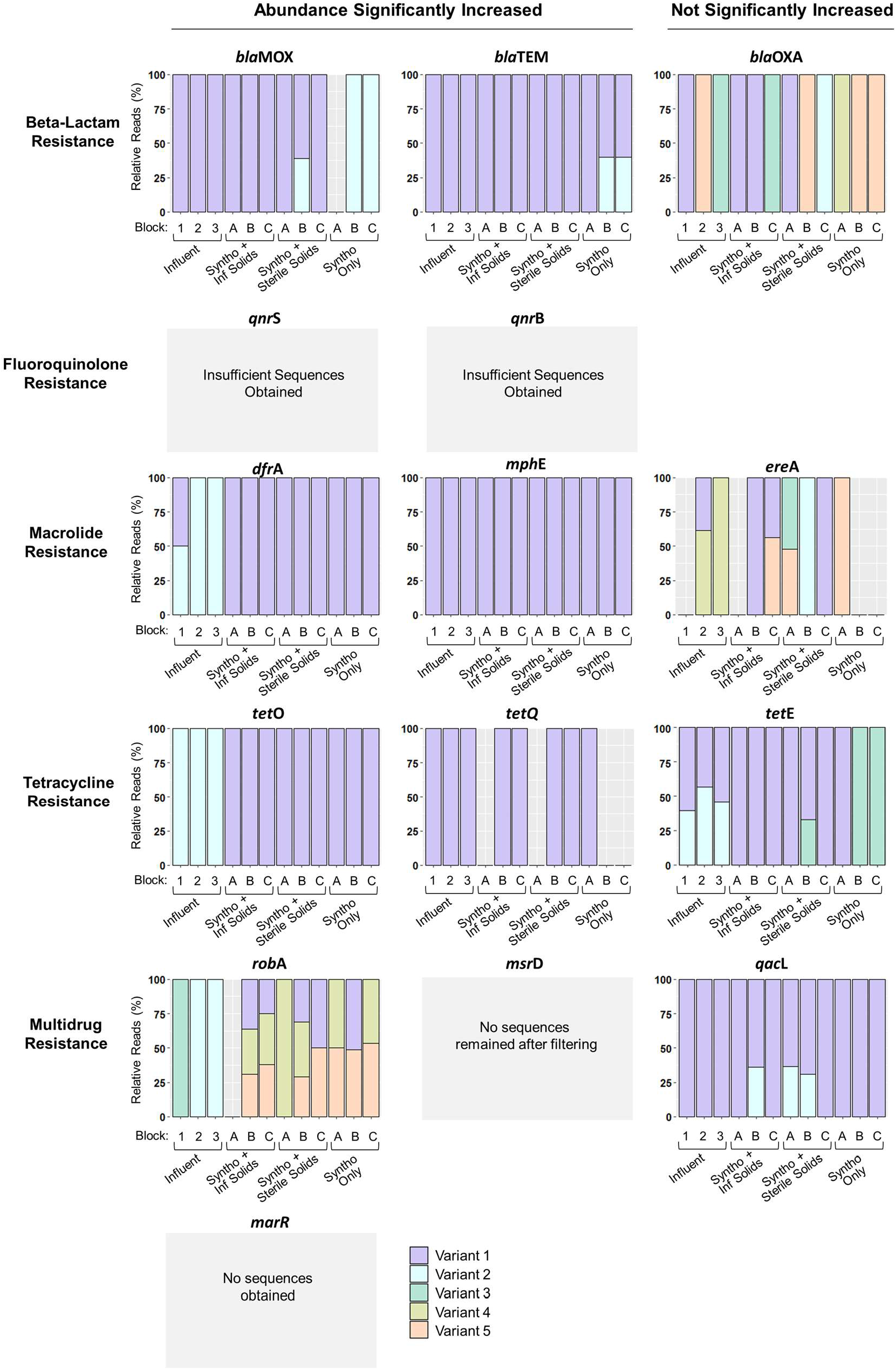
ARG sequence variants detected using multiplexed amplicon sequencing. Reactors with immigration received Syntho and Influent (Inf) Solids. Reactors receiving Syntho and Sterile Influent Solids acted as a substrate control. Reactors with Syntho only acted as a no immigration control. Each sample represents a pool of biological triplicates. Block represents the inoculum of the reactor A-La Prairie Mixed Liquor, B-Cowansville Mixed Liquor, C-Pincourt Mixed Liquor. Influent 1, 2 and 3 represent the order in which they were fed to the reactors during Phase 2, with Influent 3 received for the final solid retention time (SRT). All reactor samples analysed were obtained from the final day of Phase 2.

The concurrence profiles between the influent and the activated sludge reactors could be classified into 6 similarity groups that were typically confined either to the ARGs that significantly increased in abundance with influent immigration (Groups 1 to 3) or to ARGs with abundances that statistically remained unchanged (Groups 4 to 6; Figure 4). As summarised in Table 1, *Group 1* consists of ARGs for which the same and single variant was observed in all samples (genes *mph*E, and *tet*Q). This concurrence profile is consistent with direct immigration causing the significant increases in the abundances of these genes. *Group 2* collates ARGs with a variant observed only in the control reactor samples, but not in the live immigration reactor or the influent samples (gene *bla*MOX and *bla*TEM). This concurrence profile is also consistent with direct immigration overwhelming the detection of the variant observed in the control reactor. *Group 3* concurrence profiles are characterized by variants observed only in the influent, but not in the reactors (genes *dfr*A, *tet*O, *rob*A). This profile appears to follow a counter selection dynamic, whereby one or more of the influent sequence variants did not successfully immigrate between the influent and activated sludge. This demonstrated that the presence of a given ARG sequence variant in the influent and the significant increase of an ARG with immigration, does not definitively predict the presence of the variant within the activated sludge as could have been inferred using ddPCR results alone.

**Table 1:** Concurrence profiles of ARG sequence variants in the influent and reactor samples

| Group | Genes | Description | Conclusions |
| --- | --- | --- | --- |
| 1 | *mphE*, *tetQ* | Same single variant observed in all samples | Consistent with direct immigration |
| 2 | *bla*MOX,<br>*bla*TEM | Unique variant present in control reactors that is not detected with immigration or in the influent solids | Direct immigration and suppression dynamics |
| 3 | *dfrA*, *tetO*,<br>*robA* | Unique variants present in the influent solids only | Counter Selection |
| 4 | *tetE* | Both 2 and 3: Unique variants present in the control groups and influent solids | Mixture of direct immigration, counter selection and suppression dynamics |
| 5 | *qacL* | Unique variant only present in reactors receiving influent solids (live and autoclaved) | Possible direct immigration of variants occurring below detection limit |
| 6 | *bla*OXA,<br>*ereA* | Numerous variants distributed throughout the influent and reactor samples | Variants randomly distributed |

The *Group 4* profile which was only observed for *tet*E appeared to be a mixture of Group 2 and 3 profiles, suggesting that these two mechanisms could be at interplay. Group 3 and 4 ARGs exemplify the complexity of interactions at the interface between the influent and activated sludge. Considering the Procrustes results (Figure 3), the increase in the abundances of these ARGs could be associated with the increase in relative abundance of the bacterial populations that carry these ARGs instead (Group 3) or simultaneously (Group 4) to direct ARG immigration.

The last two profiles of concurrences were observed for ARGs that did not significantly increase in abundance (Figure 4). *Group 5* profile was only observed for *qac*L and showed variants that were only observed in some samples from reactors receiving influent solids (both live and autoclaved). These variants were in low abundances in these samples. Thus, they may have been below the detection limit in the influent or control reactors, and immigration made them detectable in some circumstances possibly due to small variations in community compositions as in Groups 3 and 4. Finally, the genes of *Group 6* (*bla*_OXA_ and *ere*A) harboured 5 variants each present seemingly at random in samples from the influent or the reactors under different experimental conditions. Consequently, no clear profiles could be identified, and immigration did not appear to affect these genes.

## 4 Discussion

### 4.1 Dynamics of ARG Sequence Variants between Influent and Activated Sludge

Droplet digital PCR revealed that the abundance of 11 of the 15 ARGs quantified increased with immigration. However, it remained to be seen whether the increase in the abundance of ARGs with immigration was due to direct ARG transport or other changes associated with influent immigration. Recent studies have demonstrated ARG sequence variants to occur in samples originating from different sources (Zhang et al., 2021). Therefore, it was hypothesised that the influent wastewater, which is strongly influenced by anthropogenic activities (Guo et al., 2019), could contain different ARG sequence variants compared to the activated sludge. The detection of unique ARG sequence variants would enable a more detailed analysis of the immigration dynamics between the influent and activated sludge.

Based upon the neutral model and the assumption of ecological equivalence (i.e., equal fitness of competitors), sequence variants detected in the influent would be expected to immigrate into the activated sludge (Harris et al., 2017). In both group 3 and 4, unique sequence variants were observed in the influent, which could have been introduced into the reactors with immigration. However, in only one instance was sequence variant immigration unequivocally observed in the current study (Figure 4). Among the *bla*MOX gene, a new sequence variant was introduced into the reactors with immigration (Figure 4). With the introduction of this sequence variant, the indigenous *bla*MOX sequence variant 2, which was present without immigration, was no longer detected, suggesting either competition among the hosts of these ARG sequence variants or simply that the low concentration of the indigenous resident variant was overwhelmed by the immigrant variant. The *bla*MOX sequence variant 2 was also detected in one of the reactors receiving sterilised influent solids, suggesting that it may have persisted after autoclaving. Counter selection of ARG sequence variants between the influent and activated sludge was more frequently observed, suggesting that ARG immigration is likely not dictated solely by neutral processes and is impacted by other factors. This conclusion is supported by recent studies which have demonstrated that in WWTP samples obtained from five different countries, the resistome composition varies between the influent and activated sludge (Dai et al., 2022).

Despite counter selection occurring, a significant increase in the abundance of many ARGs was observed in the reactors receiving active influent solids. Supported by the results from Procrustes analysis, this suggests that the abundance is impacted by the microbial community changes caused by immigration rather than direct ARG immigration between the influent and activated sludge. This is consistent with previous studies that have identified the microbial community composition to be the main driver of AMR content in numerous environments (Munck et al., 2015; Luo et al., 2017).

The results from ARG amplicon sequencing demonstrate the complexity of ARG dynamics at the interface between the influent and activated sludge. Given the frequent observation of counter selection of ARG sequence variants, PCR based approaches alone cannot be used to accurately predict the AMR the activated sludge or infer the origin of ARGs. These data also support the contrasting results reported in the literature, which are likely influenced by the methods used and targets selected. Future studies should consider the use of amplicon sequencing approaches to enable a more accurate assessment of ARG dynamics and source tracking.

### 4.2 Database Information on Genetic Context of ARG Sequence Variants

Given the differences in the environmental conditions and microbial community between the influent and mixed liquor, counter selection of ARG variants was not surprising. The successful immigration of a given ARG sequence variant is likely associated with the ability of the host to adapt and establish itself under different environmental conditions, or the mobility of the gene that may be an indicator of the likelihood of transfer to hosts better adapted to the reactor environment. To further investigate these factors, ARG sequence variants were analysed using the NCBI database and PLSDB (Galata et al., 2019) to gain information about previously reported hosts and gene occurrence on plasmids (Table S3). To exemplify this approach and establish hypotheses for future work, genes from group 2-5 (Table 1) were analysed, as unique variants were detected in either the reactors or influent samples.

In group 2, analysis of the *bla*TEM ARG using the NCBI database revealed the two variants to have previously been reported in similar bacterial hosts (Table S3), which included taxa from the genus *Acinetobacter*, *Enterobacter* and *Klebsiella* to name a few. The *bla*TEM variant 1 has been previously reported to occur on plasmids in numerous settings including in humans, animals and the environment (water) (Galata et al., 2019). Plasmids containing the *bla*TEM variant 2 have been reported in fewer settings, which included soil, feed additives and *Orcytes gigas* (rhinoceros beetle)(Galata et al., 2019). Within the influent and reactor samples *bla*TEM variant 1 was the most frequently observed. Given that both *bla*TEM variants have been previously reported among similar hosts, it could be suggested that the persistence of variant 1 is linked to the ability of plasmids containing this variant to move between environments.

In group 3, analysis of the ARG sequence variants of the *dfra*A gene found that *dfr*A sequence variant 1 corresponded to the *dfr*A-5 gene, and sequence variant 2 the *dfr*A-14 variant of the gene (Alcock et al., 2020). Sequence variant 1 (*dfr*A-5) and sequence variant 2 (*dfr*A-14) are phylogenetically closely related (Sánchez-Osuna et al., 2020) and are commonly observed within integrons and on plasmids. A 2011 review of class 1 and 2 integrons in pathogenic gram-negative bacteria identified *dfr*A-5 (sequence variant 1) in chicken and pig samples, as well as estuarine and carriage water. In the same study, the dfrA-14 gene was reported in pig, cattle and chicken samples, but not in aquatic settings (Stokes and Gillings, 2011). More recently, *dfr*A-14 has been observed in surface waters (Kohler et al., 2020) and WWTP effluent (Che et al., 2022). The widespread detection of both sequence variant 1 and 2 in clinical, agricultural, and environmental settings does not support the theory of selection due to niche differences between the influent and reactors. However, both sequence variant 1 and 2 have been observed in similar hosts, suggesting competition may occur resulting in the loss of sequence variant 2 (Table S3).

In group 4, similar selection patterns were observed with the *tet*E ARG (Figure 5c). The influent solids contained two sequence variants of the *tet*E ARG. Sequence variant 1 was detected in the reactors even without immigration, whilst sequence variant 2 did not successfully immigrate. The NCBI database returned no matches with 100% identity for sequence variant 1. The hosts of sequence variant 2 appeared to somewhat overlap with those of sequence variant 3 that was detected within the reactors. Interestingly, it was observed that sequence variant 2 was primarily reported to occur on plasmids in the NCBI database (NCBI Research Coordinators, 2013), whilst sequence variant 3 was more commonly chromosomal. Previous studies have demonstrated chromosomal mutations to carry a larger fitness cost than plasmid acquired resistance (Vogwill and Maclean, 2015), however, sequence variant 3 was found to persist in the reactors and sequence variant 2 was undetected. Sequence variant 2 has been associated with numerous plasmids found in water environments including PC1579, a conjugative plasmid recently reported to carry a novel Metallo-β-lactamase gene (Cheng et al., 2021), and plasmid pWLK-NDM, which was identified in environmental isolates carrying *bla*NDM-1 and *bla*KPC-2 resistance genes (Dang et al., 2020). Given the previous reports of sequence variant 2 in aquatic environments, it would be expected that ARB carrying this gene would be capable of growing within the reactor environment. However, sequence variant 2 remained undetected in the activated sludge suggesting again that competition for resources may be occurring.

### 4.3 Development of Multiplexed Amplicon Sequencing

Multiplexed amplicon sequencing of ARGs is a relatively new technique requiring optimisation to ensure all primer pairs work successfully in tandem (Smith et al., 2022; Gibson et al., 2023b). Overall, eleven of the fifteen ARG targets produced data on amplicon sequence variants. Of the primers not successfully producing sequence variant information, this was often due to low levels of amplification. Considering the stringent Poisson distribution-based filters applied to the amplicon sequencing data, low count sequence variants may also have been excluded during data processing. Future method developments should optimise primer design to ensure sufficient sequencing reads are obtained for all positive targets to improve resolution.

Future development of the multiplexed amplicon sequencing approach should also carefully consider which region of the ARG sequence to target. Among group 1 of concurrence profiles, low sequence diversity was found in the targeted region of the gene. The multiplexed amplicon sequence targets used herein with Illumina MiSeq protocols include only a small section of the ARG sequence (around 275 bp). Consequently, sequence diversity in other regions may have been missed. By considering regions of high genetic diversity in future design, this technique could be optimised to cover mutations of concern for example those impacting the phenotypic resistance.

### 4.4 Future Application of ARG Sequence Diversity Analysis

The monitoring of ARG sequence variants at the immigration interface revealed various immigration patterns such as (i) suppression of the indigenous activated sludge sequence variant by the immigrant, or conversely (ii) counter selection and complete immigration failure of the influent sequence variant. These immigration profiles are reported for the first time here and highlight the crucial information that can be gained using our multiplex amplicon sequencing techniques.

Unique sequence variants were observed among the influent and reactor samples, which highlights the potential for amplicon sequencing approaches to be utilised in the future for ARG source tracking purposes. To enable this, widespread sampling of reservoirs of antimicrobial resistance should be considered to identify ARG sequence variant markers present in different environments. Compared to techniques such as full metagenome sequencing, the novel amplicon sequencing approach applied here is relatively low cost and produces data that is more easily managed. In the future, such techniques could be applied for source tracking of ARG contamination in the environment and assessment of potential ARG mobility.

## 6 Author Contributions

Claire Gibson designed the study, conducted the digital droplet PCR, developed the multiplexed amplicon sequencing approach, analysed the data and prepared the manuscript. Susanne A. Kraemer developed the multiplexed amplicon sequencing approach, developed the bioinformatics pipeline, processed the sequencing data and reviewed the manuscript. Natalia Klimova developed the multiplexed amplicon sequencing approach and prepared samples for sequencing. Dominic Frigon obtained funding, supervised the research and revised the manuscript.

## 8 Acknowledgments

We would like to thank Sophie Zhang, Julia Qi, Carlos Vasquez Ochoa, Nouha Klai, Zeinab Bakhshijooybari and Shameem Jauffur for their assistance with reactor operation. We would also like to acknowledge the assistance of the operators and staff at Cowansville, La Prairie and Pincourt wastewater treatment plants for access to the facilities and help with sampling. Finally, we are indebted to Biorad for the training provided and loan of the digital droplet PCR equipment.

