## Supplementary Information for "Antibiotic Resistance Gene Variant Sequencing is Necessary to Reveal the Complex Dynamics of Immigration from Sewers to Activated Sludge"

**Table S1-** Primers used for droplet digital PCR

| Target | Forward Sequence (5'-3') | Reverse Sequence (5'-3') | Annealing Temp. (°C) | Resistance Drug Class[1] |
|---|---|---|---|---|
| *bla*MOX | ACCAGCTCGGCGGATCTG | GAGCCGGTCTTGTTGAAGAGC | 65 | Beta-Lactam |
| *bla*OXA | AGGCACGATAGTTGTGGCAGAC | GTAGAATTCCGCATTGCTGATCGC | 65 | Beta-Lactam |
| *bla*TEM | GAACCGGAGCTGAATGAAGCC | CGGGAGGGCTTACCATCTGG | 65 | Beta-Lactam |
| *dfr*A | ACATACCCTGGTCCGCGAAAG | CGCCACCAGACACTATAACGTGA | 65 | Diaminopyrimidines |
| *ere*B | CAGCTCATCGATCACCTCATGAAACCG | CACGTACGGAAGTATCTCCCTCAA | 64 | Macrolide |
| *mar*R | CACAGTTTAAGGTGCTCTGCTCTATCC | GCAAATACTCAAGTGTTGCCACTTCG | 63 | Multi-drug |
| *mph*E | AAGTGAGCAATTGGAAACCCGCTA | AGGCCGCTGCTCTTTCTAAAGTC | 65 | Macrolide |
| *msr*D | GGCAAGCTAGGTGTTGAGCAATTAG | TCCTTCACGGTCTAAATGGCTCGTA | 65 | Multi-drug |
| *qac*L | GTTGCAATCTTTGGCGAGGTCA | CGCTGACCTTGGATAGCAGGTTTAGAAC | 63 | Multi-drug |
| *qnr*B | CGACCTGAGCGGCACTGAATTTA | GCTCGCCAGTCGAAAGTCGAA | 65 | Fluoroquinolones |
| *qnr*S | ATGCCAGCTTGCGATGGCAAA | GTGGCATAAATTAGCACCCTGTAGGC | 65 | Fluoroquinolones |
| *rob*A | TCAAATGCGCGTGCAGTTCTGG | GTAGCGCTCAATATCCTGACCTTTAC | 63 | Multi-drug |
| *tet*E | TGATTGCTGGACCAGTCATTGG | CCATACGAAGCGCTCTTCTCC | 64 | Tetracycline |
| *tet*O | GCAGGGACAGAACTATTAGAGCCATATC | GCTAACTTGTGGAACATATGCCGAAC | 64 | Tetracycline |
| *tet*Q | TGGATTGAAGACCCGTCTTTGTCC | AGCAGGTGTACTTACCGGGCTATA | 65 | Tetracycline |

**Table S2-** Log change in ARG between the influent and mixed liquor

| Gene | Log Copies/16S rRNA | | Log Copies/L | | Log change |
|---|---|---|---|---|---|
| | Influent | Reactors | Influent | Reactors | |
| *bla*MOX | -2.14 + 0.31 | -3.38 + 0.13 | 7.43 | 6.91 | -0.52 |
| *bla*TEM | -2.20 + 0.06 | -3.87 + 0.13 | 7.26 | 6.41 | -0.85 |
| *bla*OXA | -1.68 + 0.41 | -2.03 + 0.29 | 7.89 | 8.31 | 0.42 |
| *qnr*S | -2.18 + 0.32 | -3.81 + 0.16 | 7.37 | 6.48 | -0.89 |
| *qnr*B | -1.86 + 0.15 | -3.67 +0.18 | 7.62 | 6.63 | -0.99 |
| *dfr*A | -2.11 + 0.10 | -3.79 + 0.16 | 7.35 | 6.50 | -0.85 |
| *mph*E | -1.34 + 0.15 | -2.39 + 0.15 | 8.16 | 7.90 | -0.26 |
| *ere*A | -2.88 + 0.01 | -4.09 + 0.05 | 6.58 | 6.18 | -0.40 |
| *tet*O | -2.88 + 0.05 | -3.86 + 0.01 | 6.58 | 6.41 | -0.17 |
| *tet*Q | -2.09 + 0.07 | -3.74 + 0.11 | 7.38 | 6.53 | -0.84 |
| *tet*E | -2.89 + 0.41 | -3.54 + 0.13 | 6.71 | 6.74 | 0.04 |
| *rob*A | -2.99 + 0.07 | -4.95 + 0.13 | 6.47 | 5.33 | -1.14 |
| *msr*D | -2.57 + 0.12 | -3.69 + 0.07 | 6.89 | 6.58 | -0.31 |
| *qac*L | -0.43 + 0.12 | -2.32 + 0.07 | 9.06 | 7.92 | -1.11 |
| *mar*R | -3.06 + 0.27 | -4.75 + 0.10 | 6.44 | 5.52 | -0.92 |

Log copies calculated by multiplying copies 16S rRNA gene/ng-DNA, Yield DNA/VSS, reactor VSS and relative ARG concentration. These values are available in Table S4 of Gibson et al., 2023.

**Table S3-** Sequence Variants Obtained from Targeted Amplicon Sequencing (Figure 4 & 5)

| Gene | Allele Sequence | NCBI BLAST/CARD Result |
|---|---|---|
| *bla*MOX<br>Variant 1 | >consensus_1<br>CTGAAGTTTGTCGGCGCCAACATG<br>ACAGGCACCGGGGACGAGGCGAT<br>GCAGCAGGCGATTGCCCTGACCCA<br>CAAGGGGGTTTACTCGGTGGGTGC<br>CATGACTCAGGGGCTCGGCTGGG<br>AGAGTTACGCCTATCCCGTGACCG<br>AAGAGACCTTGCTTGCAGGCAACT<br>CGGGCAAGGTGATCCTCGAGGCC<br>AACCCGACGGCGCCCGCCTCCAAC<br>GAGACGGGTAGCCAGGT | *Aeromonas media*<br>Uncultured bacterium |
| Variant 2 | >consensus_2<br>CTGCGCTTTGTGAAGGCCAACATC<br>AGCGGGGTGGATAATGCGGCCAT<br>GCAGCAGGCCATCGATCTGACTCA<br>CCAGGGCCAGTATGCGGTGGGGG<br>AGATGACCCAGGGACTGGGCTGG<br>GAGCGTTACCCCTATCCCGTCAGC<br>GAGCAGACGCTGCTGGCGGGCAA<br>CTCCCCGGCGATGATTTACAATGC<br>CAACCCGGCGGCGCCCGCGCCCG<br>CTGCGGCAGGGCACCCTGT | No 100% identity found<br><br>Highest percentage identity reported: 99.13 % Query Cover: 100 % |
| *tet*E<br>Variant 1 | >consensus_1<br>TGGTTTCCGATCTTGATTGCTGGA<br>CCAGTCATTGGTGGTTTTGCAGGT<br>CAACTTTCGGTACAGGCACCGTTT<br>ATGTTCGCTGCTGCCATTAACGGG<br>CTGGCATTTCTGGTCTCCCTATTCA<br>TTTTACATGAGACCCATAATGCTA<br>ATCAGGTTAGTGACGAGATAAAG<br>AATGAAACAATCAATGAAACCAC<br>ATCCTCCATACGCGAGATGATCTC<br>CCCATTATCGGGATTGCTAGTTGT<br>CTTTTTCATCATTCAATTGATTGGC<br>CAAATCCCTGCAACATTATGGGTT<br>TTATTC | No 100% identity found<br><br>Highest percentage identity reported: 100 % Query Cover: 95 % |
| Variant 2 | >consensus_2<br>TGGTTTTGCAGGTCAACTTTCGGT<br>ACAGGCACCGTTTATGTTCGCTGC<br>TGCTATTAACGGGCTGGCATTTCT<br>GGTCTCCCTATTCATTTTACATGA<br>GACCCATAATGCTAATCAGGTTAG<br>TGACGAGATAAAGAATGAAACAA<br>TCAATGAAACCACATCCTCCATAC<br>GCGAGATGATCTCCCCATTATCGG<br>GATTGCTAGTTGTCTTTTTCATCAT<br>TCAATTGATTGGCCAAATCCCTGC<br>AACATTATGGGTTTTATTC | *Aeromonas caviae*<br>*Aeromonas hydrophilia*<br>*Aeromonas media*<br>*Aeromonas salmonicida*<br>*Aeromonas veronii*<br>*Raoultella ornithinolytica*<br>*Vibrio alginolyticus*<br>*Vibrio parahaemolyticus* |

| Gene | Allele Sequence | NCBI BLAST/CARD Result |
| --- | --- | --- |
| *tet*E (continued)<br>Variant 3 | >consensus_3<br>TGGTTTTGCCGGTCAACTTTCGGT<br>ACAGGCACCGTTTATGTTCGCTGC<br>TGCTATTAACGGGCTGGCATTTCT<br>GGTCTCCCTATTCATTTTACATGA<br>GACCCATAATGCTAATCAGGTTAG<br>TGACGAGTTAAAGAATGAAACAA<br>TCAATGAAACCACATCCTCCATAC<br>GCGAGATGATCTCCCCATTATCGG<br>GATTGTTAGTTGTCTTTTTCATCAT<br>TCAATTGATTGGCCAAATCCCCGC<br>AACATTATGGGTTTTATTC | *Aeromonas caviae*<br>*Aeromonas media*<br>*Aeromonas veronii*<br>*Aeromonas sp.*<br>*Enterobacter cloacae*<br>*Yersinia ruckeri*<br>*Escherichia coli*<br>*Aeromonas hydrophila*<br>*Aeromonas dhakensis* |
| *bla*TEM<br>Variant 1 | >consensus_1<br>ATACCAAACGACGAGCGTGACAC<br>CACGATGCCTGCAGCAATGGCAA<br>CAACGTTGCGCAAACTATTAACTG<br>GCGAACTACTTACTCTAGCTTCCC<br>GGCAACAATTAATAGACTGGATG<br>GAGGCGGATAAAGTTGCAGGACC<br>ACTTCTGCGCTCGGCCCTTCCGGC<br>TGGCTGGTTTATTGCTGATAAATC<br>TGGAGCCGGTGAGCGTGGGTCTC<br>GCGGTATCATTGCAGCACTGGGG | *Acinetobacter baumannii*<br>*Acinetobacter johnsonii*<br>*Acinetobacter towneri*<br>*Aeromonas hydrophila*<br>*Aeromonas veronii*<br>*Bacillus subtilis*<br>*Bacteroides fragilis*<br>*Chlamydia trachomatis*<br>*Citrobacter amalonaticus*<br>*Citrobacter freundii*<br>*Citrobacter koseri*<br>*Citrobacter portucalensis*<br>*Citrobacter werkmanii*<br>*Citrobacter youngae*<br>*Clostridioides difficile*<br>*Cronobacter sakazakii*<br>*Enterobacter asburiae*<br>*Enterobacter chengduensis*<br>*Enterobacter cloacae*<br>*Enterobacter hormaechei*<br>*Enterobacter kobei*<br>*Enterobacter roggenkampii*<br>*Enterobacter spp.*<br>*Enterobacteriaceae*<br>*Enterococcus faecium*<br>*Escherichia albertii*<br>*Escherichia coli*<br>*Escherichia fergusonii*<br>*Escherichia marmotae*<br>*Haemophilus influenzae*<br>*Haemophilus parainfluenzae*<br>*Kingella kingaegi*<br>*Klebsiella aerogenes*<br>*Klebsiella huaxiensis*<br>*Klebsiella michiganensis*<br>*Klebsiella oxytoca*<br>*Klebsiella pneumoniae*<br>*Klebsiella quasipneumoniae*<br>*Leclercia adecarboxylata*<br>*Morganella morganii*<br>*Mycobacterium tuberculosis*<br>*Neisseria gonorrhoeae*<br>*Pasteurella multocida*<br>*Proteus mirabilis*<br>*Proteus vulgaris*<br>*Providencia rettgeri*<br>*Providencia stuartii*<br>*Pseudomonas aeruginosa*<br>*Pseudomonas putida*<br>*Raoultella planticola*<br>*Salmonella enterica*<br>*Serratia marcescens*<br>*Shigella boydii*<br>*Shigella dysenteriae*<br>*Shigella flexneri*<br>*Shigella sonnei*<br>*Staphylococcus aureus*<br>*Streptococcus suis*<br>*Vibrio cholerae*<br>*Vibrio parahaemolyticus* |

| Gene | Allele Sequence | NCBI BLAST/CARD Result | |
|---|---|---|---|
| *bla*TEM<br>(continued)<br>Variant 2 | >consensus_2<br>ATACCAAACGACGAGCGTGACAC<br>CACGATGCCTGTAGCAATGGCAAC<br>AACGTTGCGCAAACTATTAACTGG<br>CGAACTACTTACTCTAGCTTCCCG<br>GCAACAATTAATAGACTGGATGG<br>AGGCGGATAAAGTTGCAGGACCA<br>CTTCTGCGCTCGGCCCTTCCGGCT<br>GGCTGGTTTATTGCTGATAAATCT<br>GGAGCCGGTGAGCGTGGGTCTCG<br>CGGTATCATTGCAGCACTGGGG | *Acinetobacter baumannii*<br>*Acinetobacter haemolyticus*<br>*Bacillus cereus*<br>*Bacillus subtilis*<br>*Bacillus velezensis*<br>*Bifidobacterium longum*<br>*Burkholderia cepacia*<br>*Burkholderia lata*<br>*Chlamydia trachomatis*<br>*Clostridioides difficile*<br>*Clostridium botulinum*<br>*Cronobacter sakazakii*<br>*Enterobacter cloaceae*<br>*Enterococcus faecium*<br>*Escherichia coli*<br>*Faecalibacterium prausnitzii*<br>*Helicobacter pylori*<br>*Klebsiella michiganensis*<br>*Klebsiella pneumoniae*<br>*Klebsiella quasipneumoniae* | *Leclercia adecarboxylata*<br>*Legionella pneumophila*<br>*Mycobacterium tuberculosis*<br>*Mycoplasma mycoides*<br>*Neisseria meningitidis*<br>*Propionibacterium freudenre*<br>*Pseudomonas aeruginosa*<br>*Rhizobium leguminosarum*<br>*Ruthenibacterium lactatiform*<br>*Salmonella enterica*<br>*Staphylococcus aureus*<br>*Staphylococcus hominis*<br>*Staphylococcus saprophyticus*<br>*Streptococcus agalactiae*<br>*Streptococcus lutetiensis*<br>*Streptococcus pneumoniae*<br>*Vibrio parahaemolyticus* |
| *dfr*A<br>Variant 1 | >consensus_1<br>GGAGAGCAGCTACTCTTTAAAGCC<br>TTGACGTACAACCAGTGGCTTTTG<br>GTGGGCCGCAAGACGTTCGAATCT<br>ATGGGAGCACTCCCTAATAGGAA<br>ATACGCGGTCGTTACTCGCTCAGC<br>CTGGACGGCCGATAATGACAACG<br>TAATAGTATTCCCGTCGATCGAAG<br>AGGCCATGTACGGGCTGGCTGAA<br>CTCACCGA | *Acinetobacter baumannii*<br>*Aeromonas caviae*<br>*Citrobacter koseri*<br>*Comamonas testosteroni*<br>*Enterobacter hormaechei*<br>*Enterobacter roggenkampii*<br>*Escherichia albertii*<br>*Escherichia coli*<br>*Escherichia fergusonii*<br>*Klebsiella michiganensis*<br>*Klebsiella pneumoniae*<br>*Klebsiella quasipneumoniae* | *Proteus mirabilis*<br>*Proteus vulgaris*<br>*Vibrio cholerae*<br>*Pseudomonas aeruginosa*<br>*Salmonella enterica*<br>*Serratia marcescens*<br>*Shigella boydii*<br>*Shigella flexneri*<br>*Shigella sonnei* |
| Variant 2 | >consensus_2<br>GGGGAGCAGCTACTTTTTAAAGCA<br>TTGACCTACAATCAGTGGCTTCTG<br>GTGGGTCGCAAGACGTTTGAATCT<br>ATGGGCGCACTCCCCAATAGGAA<br>ATACGCGGTCGTTACCCGCTCAGG<br>TTGGACATCAAATGATGACAATGT<br>AGTTGTATTTCAGTCAATCGAAGA<br>GGCCATGGACAGGCTAGCTGAATT<br>CACCGG | *Aeromonas hydrophila*<br>*Aeromonas veronii*<br>*Citrobacter amalonaticus*<br>*Citrobacter freundii*<br>*Citrobacter portucalensis*<br>*Citrobacter werkmanii*<br>*Citrobacter youngae*<br>*Enterobacter asburiae*<br>*Enterobacter cloacae*<br>*Enterobacter hormaechei*<br>*Enterobacter kobei*<br>*Enterobacter roggenkampii*<br>*Escherichia albertii*<br>*Escherichia coli*<br>*Escherichia fergusonii*<br>*Klebsiella michiganensis*<br>*Klebsiella grimontii*<br>*Klebsiella pneumoniae*<br>*Klebsiella quasipneumoniae* | *Morganella morganii*<br>*Proteus mirabilis*<br>*Proteus vulgaris*<br>*Providencia rettgeri*<br>*Providencia stuartii*<br>*Pseudomonas aeruginosa*<br>*Raoultella planticola*<br>*Salmonella enterica*<br>*Serratia marcescens*<br>*Shewanella putrefaciens*<br>*Shigella boydii*<br>*Shigella dysenteriae*<br>*Shigella flexneri*<br>*Shigella sonnei*<br>*Vibrio cholerae* |

| Gene | Allele Sequence | NCBI BLAST/CARD Result | |
|---|---|---|---|
| *bla*OXA<br>Variant 1 | >consensus_1<br>GAACGCCAAGCGGATCGTGCCAT<br>GTTGGTTTTTGATCCTGTGCGATC<br>GAAGAAACGCTACTCGCCTGCATC<br>GACATTCAAGATACCTCATACACT<br>TTTTGCACTTGATGCAGGCGCTGT<br>TCGTGATGAGTTCCAGATTTTTCG<br>ATGGGACGGCGTTAACAGGGGCT<br>TTGCAGGCCACAATCAAGACCAA<br>GATTTGCGATCAGCAATGCGGAAT<br>TCTACAGATCGGAAGAGCACACG<br>TCT | *Acinetobacter baumannii*<br>*Aeromonas caviae*<br>*Alcaligenes faecalis*<br>*Burkholderia cenocepacia*<br>*Citrobacter freundii*<br>*Citrobacter koseri*<br>*Citrobacter portucalensis*<br>*Citrobacter werkmanii*<br>*Enterobacter asburiae*<br>*Enterobacter cloacae*<br>*Enterobacter hormaechei*<br>*Enterobacter kobei*<br>*Enterobacter roggenkampii*<br>*Escherichia coli*<br>*Klebsiella michiganensis*<br>*Klebsiella oxytoca* | *Klebsiella pneumoniae*<br>*Klebsiella quasipneumoniae*<br>*Pasteurella multocida*<br>*Proteus mirabilis*<br>*Providencia rettgeri*<br>*Providencia stuartii*<br>*Pseudomonas aeruginosa*<br>*Pseudomonas monteilii*<br>*Pseudomonas putida*<br>*Pseudomonas stutzeri*<br>*Salmonella enterica*<br>*Serratia marcescens*<br>*Shigella sonnei*<br>*Stenotrophomonas maltophili*<br>*Vibrio cholerae* |
| Variant 2 | >consensus_2<br>GAACGCCAAGCGGATCGTGCCAT<br>GTTGGTTTTTGATCCTGTGCGATC<br>GAAGAAACGCTACTCGCCTGCATC<br>GACATTCAAGATACCTCATACACT<br>TTTTGCACTTGATGCAGGCGCTGT<br>TCGTGATGAGTTCCAGATTTTTCG<br>ATGGGACGGCGTTAACAGGGGCT<br>TTGCAGGCCACAATCAAGACCAA<br>GATTTGCGACCAGCAATGCGGAAT<br>TCTACAGATCGGAAGAGCACAC | No 100% identity found<br><br>Highest percentage identity reported: 99.54 % Query Cover: 92 % | |
| Variant 3 | >consensus_3<br>GAACGCCAAGCGGATCGTGCCAT<br>GTTGGTTTTTGATCCTGTGCGATC<br>GAAGAAACGCTACTCGCCTGCATC<br>GACATTCAAGATACCTCATACACT<br>TTTTGCACTTGATGCAGGCGCTGT<br>TCGTGATGAGTTCCAGATTTTTCG<br>ATGGGACGGCGTTAACAGGGGCT<br>TTGCAGGCCACAATCAAGACCAA<br>GATTTGCGATCAGCAATGCGGAAT<br>TTAC | No 100% identity found<br><br>Highest percentage identity reported: 99.54 % Query Cover: 100 % | |
| Variant 4 | >consensus_4<br>GAACGCCAAGCGGATCGTGCCAT<br>GTTGGTTTTTGATCCTGTGCGATC<br>GAAGAAACGCTACTCGCCTGCATC<br>GACATTCAAGATACCTCATACACT<br>TTTTGCACTTGATGCAGGCGCTGT<br>TCGTGATGAGTTCCAGATTTTTCG<br>ATGGGACGGCGTTAACAGGGGCT<br>TTGCAGGCCACAATCAAGACCAA<br>GATTTGCGATCAGCGAT | *Pseudomonas aeruginosa*<br>*Pseudomonas putida*<br>*Stenotrophomonas maltophilia* | |

| Gene | Allele Sequence | NCBI BLAST/CARD Result |
|---|---|---|
| *bla*OXA (continued)<br>Variant 5 | >consensus_5<br>GAACGCCAAGCGGATCGTGCCAT<br>GTTGGTTTTTGATCCTGTGCGATC<br>GAAGAAACGCTACTCGCCTGCATC<br>GACATTCAAGATACCTCATACACT<br>TTTTGCACTTGATGCAGGCGCTGT<br>TCGTGATGAGTTCCAGATTTTTCG<br>ATGGGACGGCGTTAACAGGGGCT<br>TTGCAGGCCACAATCAAGACCAA<br>GATTTT | *Acinetobacter baumannii*<br>*Aeromonas caviae*<br>*Alcaligenes faecalis*<br>*Burkholderia cenocepacia*<br>*Citrobacter freundii*<br>*Citrobacter koseri*<br>*Citrobacter portucalensis*<br>*Citrobacter werkmanii*<br>*Enterobacter asburiae*<br>*Enterobacter cloacae*<br>*Enterobacter hormaechei*<br>*Enterobacter kobei*<br>*Enterobacter roggenkampii*<br>*Escherichia coli*<br>*Klebsiella michiganensis*<br>*Klebsiella oxytoca*<br>*Klebsiella pneumoniae*<br>*Klebsiella quasipneumoniae*<br>*Pasteurella multocida*<br>*Proteus mirabilis*<br>*Providencia rettgeri*<br>*Providencia stuartii*<br>*Pseudomonas aeruginosa*<br>*Pseudomonas monteilii*<br>*Pseudomonas putida*<br>*Salmonella enterica*<br>*Serratia marcescens*<br>*Shigella sonnei*<br>*Stenotrophomonas maltophili*<br>*Vibrio cholerae* |
| *qac*L<br>Variant 1 | >consensus_1<br>CACGACGCTCTTCCGATCTCTGTT<br>TCAATCTTTGGCGCGGTCATCGCA<br>ACTTCCGCACTGAAGTCTAGCCAT<br>GGATTCACTAGGTTAGTTCCTTCC<br>GTTGTAGTTGTGGCTGGCTACGGG<br>CTTGCGTTCTATTTCTTGTCTCTCG<br>CGCTCAAGTCCATTCCGGTCGGTA<br>TTGCTTACGCTGTATGGGCTGGGC<br>TTGGCATCGTGCTTGTGGCAGCTA<br>TTGCTTGGATTTTCCATGGCCAAA<br>AACTAGACTTCTGGGCGTTCATTG<br>GCATGGGACTTAT | No 100% identity found<br><br>Highest percentage identity reported: 99.61 % Query Cover: 93 % |
| Variant 2 | >consensus_2<br>TCGCAACTTCCGCACTGAAGTCTA<br>GCCATGGATTCACTAGGTTAGTTC<br>CTTCCGTTGTAGTTGTGGCTGGCT<br>ACGGGCTTGCGTTCTATTTCTTGTC<br>TCTCGCGGTCAAGTCCATTCCGGT<br>CGGTATTGCTTACGCTGTATGGGC<br>TGGGCTTGGCATCGTGCTTGTGGC<br>AGCTATTGCTTGGATTTTCCATGG<br>CCAAAAACTAGACTTCTGGGCGTT<br>CATTGGCATGGGACTTAT | Uncultured prokaryote |

| Gene | Allele Sequence | NCBI BLAST/CARD Result | |
|---|---|---|---|
| *mar*R<br>Variant 1 | >consensus_8<br>GCTGTGCGGGCTGCATTACCCCGG<br>TTGAACTGAAAAAAGTGTTGTCTG<br>TCGATCTCGGCGCCTTAACGCGCA<br>TGCTCGAGCGTCTGGTCTGCAAAG<br>GCTGGATTGACAGACTGCCTAACC<br>CACATGACAAACGCGGTGTGCTG<br>GTGAAACTCACCGAACACGGCGC<br>GGCAATTTGTGAGCAATGTCATCA<br>ATTAGTAGGACAAGACCTGCACC<br>AGGAATTAACAAAAAACTTAACG<br>GCGGA | *Citrobacter*<br>*Citrobacter sp.*<br>*Citrobacter amalonaticus* | |
| *tet*Q<br>Variant 1 | >consensus_1<br>CTTTTCCATAAACTCATATAGTGA<br>TGAATTGGAAATCTCGTTATATGG<br>TTTGACCCAAAAGGAAATCATACA<br>GACATTGCTGGAAGAACGATTTTC<br>CGTAAAGGTCCATTTTGATGAGAT<br>CAAGACTATCTACAAAGAACGAC<br>CTATAAAAAAGGTCAATAAGATT<br>ATTCAGATCGAAGTACCACCCAAC<br>CCTTACTGGGCCACAATAGGGCTG<br>ACTCTTGAACCCTTACCGTTAGGG<br>GCAGGGTTGCAAATCGAAAGTGA<br>CATCTCCTATGGTTATCTGAACCA<br>TTCTTTTCAAAATGCCGTTTTTGAA<br>GGGATTCGTATGTCTTGCCAATCT<br>GGTTTACATGGATGGGAAGTGAC<br>AGATCTGAAAGTAACTTTTACTCA<br>AGCCGAGTAT | *Alistipes communis*<br>*Alistipes onderdonkii*<br>*Bacteroides caccae*<br>*Bacteroides caecimuris*<br>*Bacteroides cellulosilyticus*<br>*Bacteroides dorei*<br>*Bacteroides eggerthii*<br>*Bacteroides fragilis*<br>*Bacteroides ovatus*<br>*Bacteroides salyersiae*<br>*Bacteroides sp.*<br>*Bacteroides thetaiotaomicron*<br>*Bacteroides uniformis*<br>*Bactroides xylanisolvens*<br>*Bacteroides zhangwenhongii*<br>*Butyricimonas faecalis*<br>*Coprobacter secundus*<br>*Odoribacteraceae bacterium* | *Parabacteroides distasonis*<br>*Parabacteroides goldsteinii*<br>*Parabacteroides johnsonii*<br>*Parabacteroides merdae*<br>*Parabacteroides sp.*<br>*Paraprevotella xylaniphila*<br>*Phocaeicola dorei*<br>*Phocaeicola vulgatus*<br>*Prevotella buccalis*<br>*Prevotella intermedia*<br>*Prevotella melaninogenica*<br>*Prevotella ruminocola*<br>*Prevotella sp.*<br>*Pseudoprevotella muciniphila*<br>*Riemerella anatipestifer*<br>*Sodaliphilus pleomorphus*<br>*Uncultured bacterium* |
| *mph*E<br>Variant 1 | >consensus_2<br>CAGAAAATGGTTGGATAATGATGT<br>TCTATGGGCAGATTTCACCCAATT<br>TATACATGGCGATTTATATGCTGG<br>GCATGTACTAGCTTCAAAGGATGG<br>AGCTGTTTCAGGCGTTATTGATTG<br>GTCAACAGCCCATATAGATGACCC<br>AGCGATTGATTTTGCT<br>GGGCATGTAACTTTGTTTGGAGAA<br>GAAAGCCTCAAAACTCTAATCATC<br>GAGTATGAAAAACTAGGGGGTAA<br>AGTTTGGAATAAACTATATGAACA<br>GACTTTAGAAAGAGCAGCGGCCT<br>AGATCGGAA | No 100 % identity found<br><br>Highest percentage identity reported: 100 % Query Cover: 96 % | |

| Gene | Allele Sequence | NCBI BLAST/CARD Result |
|---|---|---|
| *ere*A<br>Variant 1 | >consensus_1<br>CACGTTGATATGCTGACTCACTTG<br>TTGGCGTCCATTGATGGCCAGTCG<br>GCGGTTATTTCATCGGCAAAATGG<br>GGGGAGCTAGAAACGGCTCGGCA<br>GGAGAAAGCTATCTCAGGGGTAA<br>CCAGATTGAAGCTCCGCTTGGCGT<br>CGCTTGCCCCTGTCCTGAAAAAAC<br>ACGTCAACAGCGATTTGTTCCGAA<br>AAGCCTCTGATCGAATAGAGTCGA<br>TAGAGTATACGTTGGAAACCTTGC<br>GTATAATGAAAACTTTCTTCGATG<br>GTACCTCTC | *Providencia huaxiensis*<br>*Pandoraea sp.*<br>*Comamonas sp.*<br>*Vogesella fluminis*<br>*Uncultured bacterium*<br>*Vogesella perlucida*<br>*Comamonas koreensis* |
| Variant 2 | >consensus_2<br>CACGTTGATATGCTGACTCACTTG<br>TTGGCGTCCATTGATGGCCAGTCG<br>GCGGTTATTTCATCGGCAAAATGG<br>GGGGAGCTAGAAACGGCTCGGCA<br>GGAGAAAGCTATCTCAGGGGTAA<br>CCAGATTGAAGCTCCGCTTGGCAT<br>CGCTTGCCCCTGTCCTGAAAAAAC<br>ACGTCAACAGCGATTTGTTCCGAA<br>AAGCCTCTGATCGAATAGAGTCGA<br>TAGAGTATACGTTGGAAACCTTGC<br>GTATAATGAAAACTTTCTTCGATG<br>GTACCTCTC | No 100% identity found<br><br>Highest percentage identity reported: 99.63 % Query Cover: 100 % |
| Variant 3 | >consensus_3<br>CACGTTGATATGTTGACTCACTTG<br>TTGGCGTCCATTGATGGCCAGTCG<br>GCGGTTATTTCATCGGCAAAATGG<br>GGGGAGCTAGAAACGGCTCGGCA<br>GGAGAAAGCTATCTCAGGGGTAA<br>CCAGATTGAAGCTCCGCTTGGCGT<br>CGCTTGCCCCCGTCCTGAAAAAAC<br>ACGTCAACAGCGATTTGTTCCGAA<br>AAGCCTCTGATCGAATAGAGTCGA<br>TAGAGTATACGTTGGAAACCTTGC<br>GTATAATGAAAACTTTCTTCGATG<br>GTACCTCTC | *Achromobacter denitrificans*<br>*Enterobacter hormaechei*<br>*Escherichia coli*<br>*Helicobacter pylori*<br>*Klebsiella pneumoniae*<br>*Pseudomonas aeruginosa*<br>*Salmonella enterica* |
| Variant 4 | >consensus_4<br>CACGTTGATATGCTGACTCACTTG<br>TTGGCGTCCATTGATGGCCAGTCG<br>GCGGTTATTTCATCGGCAAAATGG<br>GGGGAGCTAGAAACGGCTCGGCA<br>GGAGAAAGCTATCTCAGGGGTAA<br>CCAGATTGAAGCTCCGCTTGGCGT<br>CGCTTGCCCCTGTACTGAAAAAAC<br>ACGTCAACAGCGATTTGTTCCGAA<br>AAGCCTCTGATCGAATAGAGTCGA<br>TAGAGTATACGTTGGAAACCTTGC<br>GTATAATGAAAACTTTCTTCGATG<br>GTACCTCTC | *Escherichia coli*<br>*Pseudomonas aeruginosa*<br>*Salmonella enterica*<br>*Serratia marcescens*<br>*Thauera humireducens*<br>*Uncultured bacterium* |

| Gene | Allele Sequence | NCBI BLAST/CARD Result |
|---|---|---|
| *ere*A | | |
| (continued) | >consensus_5 | |
| Variant 5 | CACGTTGATATGCTGACTCACTT | *Aeromonas hydrophila* |
| | GTTGGCGTCCATTGATGGCCAGT | *Escherichia coli* |
| | CGGCGGTTATTTCATCGGCAAAA | *Klebsiella pneumoniae* |
| | TGGGGGGAGCTAGAAACGGCTC | *Klebsiella oxytoca* |
| | GGCAGGAGAAAGCTATCTCAGG | *Laribacter hongkongensis* |
| | GGTAACCAGATTGAAGCTCCGCT | *Proteus mirabilis* |
| | TGGCGTCGCTTGCCCCTGTACTG | *Proteus terrae* |
| | AAAAAACACGTCAACAGCGATT | *Proteus vulgaris* |
| | TGTTCCGAAAAGCCTCTGATCGA | *Providencia rettgeri* |
| | ATAGAATCGATAGAGTATACGTT | *Pseudoalteromonas sp.* |
| | GGAAACCTTGCGTATAATGAAA | *Salmonella enterica* |
| | ACTTTCTTCGATGGTACCTCTC | *Vibrio alginolyticus* |
| | | *Vogesella perlucida* |
| *rob*A | >consensus_1 | |
| Variant 1 | CACGATTTTCTCGGCAACGCGCC | *Escherichia coli* |
| | GACCATTCCGCCAGTGCTCTACG | *Shigella dysenteriae* |
| | GCCTGAATGAAACGCGTCCGAG | *Shigella flexneri* |
| | TCAGGATAAAGACGACGAACAA | |
| | GAGGTATTCTATACCACCGCGTT | |
| | AGCCCAGGATCAGGCAGATGGC | |
| | TATGTACTGACGGGGCATCCGGT | |
| | GATGCTGCAGGGCGGCGAATAT | |
| | GTGATGTTTACCTATGAAGGTCT | |
| | GGGAACCGGCGTGCAGGAGTTT | |
| | ATCCTGACGGTATACGGAACGT | |
| | GCATGCCAATGCTCAACCTGACG | |
| | CGCC | |
| Variant 2 | >consensus_2 | |
| | CGCGACTTCCTGAGCCATGCCCC | *Enterobacter cloacae* |
| | GGCGATCCCGCCTATCCTGTATG | |
| | GTCTCAACGAAACGCATCCAAG | |
| | CCAGGAAAAAGACGACGAGCAG | |
| | GAGGTGTTCTACACCACCGCGTT | |
| | AACGCCGGAAATGGCCAATGGC | |
| | TATATTCAGGGCTCTAAACCGGT | |
| | CGTGCTGGAAGGCGGAGAGTAC | |
| | GTGATGTTCTCCTACGAAGGGCT | |
| | GGGAACGGGCGTACAGGAATTC | |
| | ATCCTGACCGTTTACGGAACATG | |
| | CATGCCTATGCTGAACCTGAATC | |
| | GCC | |

| Gene | Allele Sequence | NCBI BLAST/CARD Result |
| --- | --- | --- |
| *rob*A<br>(continued)<br>Variant 3 | >consensus_3<br>CGCGACTTCCTGAGCCACGCACC<br>GGCGATCCCGCCTATTCTGTATG<br>GTCTCAACGAAACGCACCCGAG<br>CCAGGAAAAGGA<br>CGACGAGCAGGAGGTGTTCTAC<br>ACCACCGCGCTGACGCCAGAGA<br>TGGCCAATGGCTACATTCAGGGT<br>TCAAAACCTGTCG<br>TGCTGGAAGGCGGTGAATACGT<br>GATGTTCGCCTATGAAGGGCTGG<br>GAACGGGCGTTCAGGAGTTCAT<br>CCTGACCGTTTAC<br>GGAACCTGCATGCCGATGCTGA<br>ATCTGAATCGCC | *Enterobacter cloacae* |
| Variant 4 | >consensus_4<br>CACGATTTTCTCGGCAACGCGCC<br>GACCATTCCGCCAGTGCTCTACG<br>GCCTGAATGAAACGCGTCCGAG<br>TCAGGATAAAGA<br>CGACGAACAAGAGGTATTCTAT<br>ACCACTGCGTTAGCCCAGGATCA<br>GGCAGATGGCTATGTACTGACG<br>GGGCATCCGGTGA<br>TGCTGCAGGGCGGCGAATATGT<br>GATGTTTACCTATGAAGGTCTGG<br>GAACCGGCGTGCAGGAGTTTAT<br>CCTGACAGTATAC<br>GGAACGTGCATGCCAATGCTCA<br>ATCTGACGCGCC | *Escherichia coli* |
| Variant 5 | >consensus_5<br>GCACGATTTTCTCGGCAACGCGC<br>CGACCATTCCGCCGGTGCTCTAC<br>GGCCTGAACGAAACGCGTCCGA<br>GTCAGGATAAAG<br>ACGACGAACAAGAGGTATTCTA<br>TACCACCGCGTTAGCCCAGGATC<br>AGGCAGATGGCTATGTACTGAC<br>GGGGCATCCGGTG<br>ATGCTGCAGGGCGGCGAATATG<br>TGATGTTTACCTATGAAGGTCTG<br>GGAACCGGCGTGCAGGAGTTTA<br>TCCTGACGGTATA<br>CGGAACGTGCATGCCAATGCTC<br>AACCTGACGCGCC | *Escherichia coli* |

| Gene | Allele Sequence | NCBI BLAST/CARD Result |
| --- | --- | --- |
| *tet*O<br>Variant 1 | >consensus_1<br>TCCACTTTGAAATTTATGCACCGCAGGA<br>ATATCTCTCACGGGCGTATCATGATGCT<br>CCAAGGTATTGTGCAGATATTGTA<br>AGTACTCAGATAAAGAATGACGAGGTCA<br>TTCTGAAAGGAGAAATCCCTGCTAGATG<br>TATTCAAGAATACAGGAACGATTT<br>AACTAATTTCACAAATGGGCAGGGAGTC<br>TGCTTGACAGAGTTAAAAGGATACCAGC<br>CAGCTATTGGTAAATTTATTTGCC<br>AACCCCGCCGCCCGAATAGCCGTATAGA<br>TAAG | *Bifidobacterium thermophilum*<br>*Campylobacter jejuni*<br>*Lactobacillus johnsonii*<br>*Riemerella anatipestifer*<br>*Streptococcus galloyticus*<br>*Streptococcus phage*<br>*Streptococcus suis*<br>*Uncultured bacterium* |
| Variant 2 | >consensus_2<br>TCCACTTTGAAATTTATGCACCGCAGGA<br>ATATCTCTCACGGGCGTATCATGATGCT<br>CCAAGGTATTGTGCAGATATTGTA<br>AGTACTCAGATAAAGAATGACGAGGTCA<br>TTCTGAAAGGAGAAATCCCTGCTAGATG<br>TATTCAAGAATACAGGAACGATTT<br>AACTTATTTCACAAATGGGCAGGGAGTC<br>TGCTTGACAGAGTTAAAAGGATACCAGC<br>CAGCTATTGGTAAATTTATTTGCC<br>AACCCCGCCGCCCGAATAGCCGTATAGA<br>TAAG | *Actinobacillus pleuropneumoniae*<br>*Anaerostipes rhamnosivorans*<br>*Bifidobacterium breve*<br>*Bifidobacterium pseudocatenulatum*<br>*Blautia massiliensis*<br>*Blautia sp.*<br>*Campylobacter coli*<br>*Campylobacter jejuni*<br>*Clostridiales genomosp,*<br>*Clostridioides difficile*<br>*Coprococcus comes*<br>*Eggerthella lenta*<br>*Emergencia timonensis*<br>*Enterocloster bolteae*<br>*Enterocloster clostridioformis*<br>*Enterococcus cecorum*<br>*Enterococcus faecalis*<br>*Enterococcus gallinarum*<br>*Erysipelotrichaceae bacterium*<br>*Eubacterium sp.*<br>*Faecalibacterium prausnitzii*<br>*Glaesserella parasuis*<br>*Lachnospiraceae bacterium*<br>*Roseburia intestinalis*<br>*Streptococcus dysgalactiae*<br>*Streptococcus porcinus*<br>*Streptococcus pyogenes*<br>*Streptococcus suis*<br>*Uncultured Blautia sp.*<br>*Uncultured Dorea sp.*<br>*Uncultured Eubacteriales bacterium*<br>*Clostridium hylemonae*<br>*Clostridium innocuum*<br>*Clostridium scindens*<br>*Clostridium symbiosum* |
